# EnteroBase in 2025: exploring the genomic epidemiology of bacterial pathogens

**DOI:** 10.1101/2024.08.15.608032

**Authors:** Nigel P. Dyer, Birgitta Päuker, Laura Baxter, Anshul Gupta, Boyke Bunk, Jörg Overmann, Margo Diricks, Viola Dreyer, Stefan Niemann, Kathryn E. Holt, Mohammed Rahman, Paul E. Brown, Richard Stark, Zhemin Zhou, Sascha Ott, Ulrich Nübel

**Affiliations:** Bioinformatics & Digital Health Services, Research Technology Platforms, University of Warwick, Coventry, UK; Leibniz Institute DSMZ, Germany - German Collection of Microorganisms and Cell Cultures, Braunschweig, Germany; German Center for Infection Research (DZIF), partner site Braunschweig-Hannover, Braunschweig, Germany; Technical University of Braunschweig, Institute of Microbiology, Braunschweig, Germany; Research Center Borstel, Leibniz Lung Center, Borstel, Germany; German Center for Infection Research (DZIF), partner site Hamburg-Lübeck-Borstel-Riems, Borstel, Germany; National and WHO Supranational Reference Laboratory for Mycobacteria, Research Center Borstel, Germany; Department of Infection Biology, Faculty of Infectious and Tropical Diseases, London School of Hygiene & Tropical Medicine, London, UK; Department of Infectious Diseases, School of Translational Medicine, Monash University, Melbourne, Victoria, Australia; Soochow University, China

## Abstract

This paper presents an update on the content, accessibility and analytical tools of the EnteroBase platform for web-based pathogen genome analysis. EnteroBase provides manually curated databases of genome sequence data and associated metadata from currently >1.1 million bacterial isolates, more recently including *Streptococcus* spp. and *Mycobacterium tuberculosis*. We have implemented the genome-based detection of antimicrobial resistance determinants and the new bubble plot graphical tool for visualising bacterial genomic population structures, based on pre-computed hierarchical clusters. Access to data and analysis tools is provided through an enhanced graphical user interface and a new application programming interface (RESTful API). EnteroBase is now being developed and operated by an international consortium, to accelerate the development of the platform and ensure the longevity of the resources built. EnteroBase can be accessed at https://enterobase.warwick.ac.uk as well as https://enterobase.dsmz.de.

**GRAPHICAL ABSTRACT:** 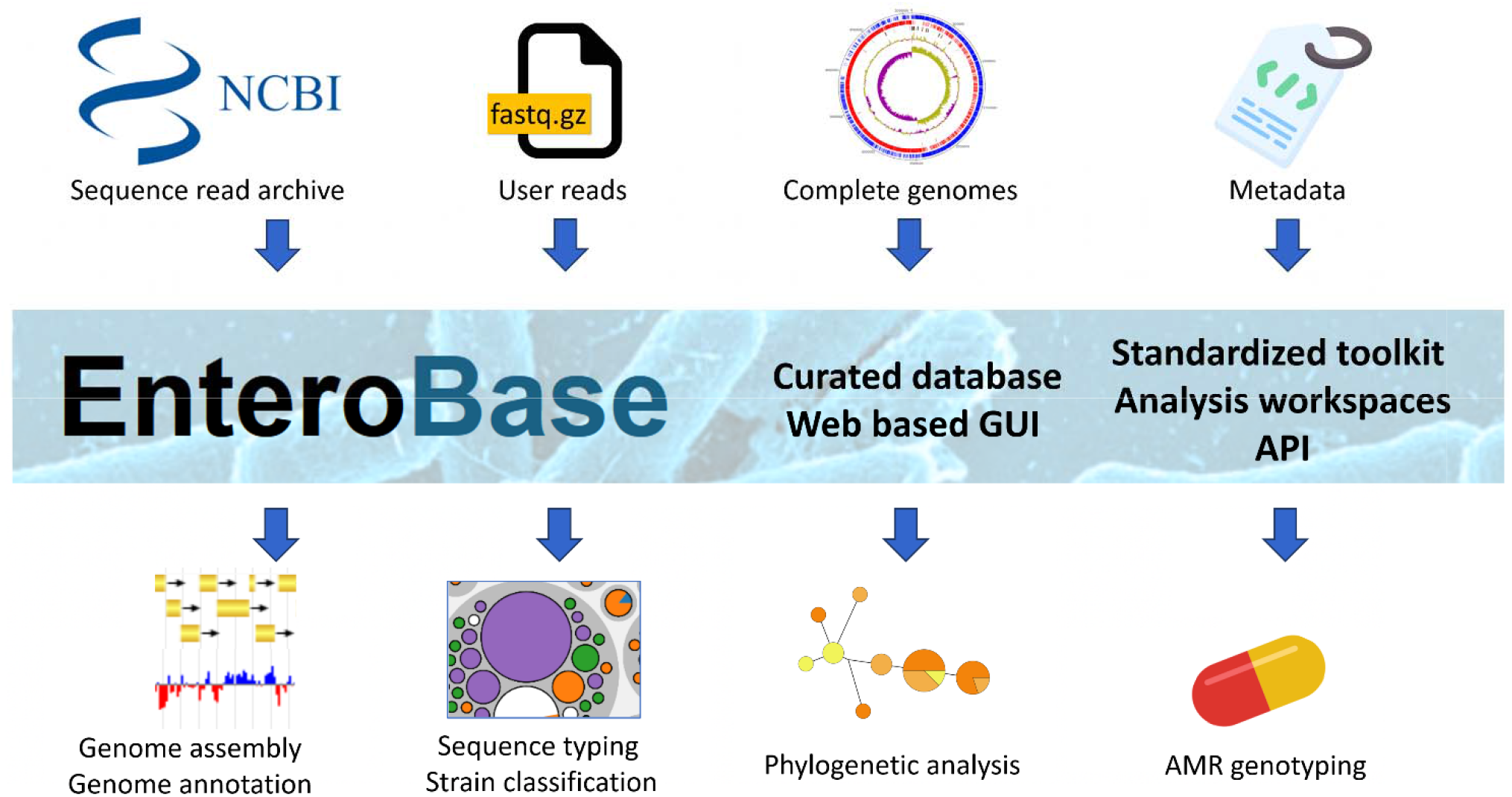

## INTRODUCTION

EnteroBase is a publicly accessible, free to use, web-based platform for pathogen genomic analyses, currently providing genome sequence data from 1,144,499 bacterial isolates (as of 31 July 2024) in a uniformly assembled and annotated format. EnteroBase is updated daily by automatically scanning the NCBI short-read archive for newly published sequence data. Users can also upload sequencing data and corresponding metadata directly. Genome assemblies that pass quality control are genotyped by using core-genome multilocus sequence typing (cgMLST) followed by a unique, multi-level hierarchical clustering approach. Genome data are presented together with associated metadata, including geographic origin, isolation date, source sample, type of disease etc. Expert microbiologists with specialised knowledge of specific pathogens actively curate the isolate-associated metadata based on scientific literature and on interaction with the scientists who submitted the sequence data. By providing users with tools for phylogenetic, epidemiological, and comparative genomic analyses, augmented by powerful graphical visualisation and search capabilities, EnteroBase serves a broad user community, ranging from medical microbiologists to clinicians, epidemiologists, population geneticists and bioinformaticians.

EnteroBase was founded in 2014, and individual databases (1,2), implemented tools (3,4), and the results obtained with them (5-7) were reported previously. The last major overview of EnteroBase was published in December 2019 (8). Since then, a number of advances have been made. In this paper, we outline selected improvements to EnteroBase content, accessibility and analytical tools.

The EnteroBase platform is now operated and developed in an international consortium including the University of Warwick (UK) and the Leibniz Institute DSMZ (Germany), accelerating the development of the platform, increasing its data security, and ensuring the longevity of the resources built. Accordingly, EnteroBase can be accessed at https://enterobase.warwick.ac.uk as well as https://enterobase.dsmz.de. The number of EnteroBase users and analysis jobs has increased steadily over recent years, and over the last 12 months, EnteroBase has been accessed by users in 151 countries worldwide, indicating the platform is widely and routinely used in many countries (Figure 1).

**Figure 1.**
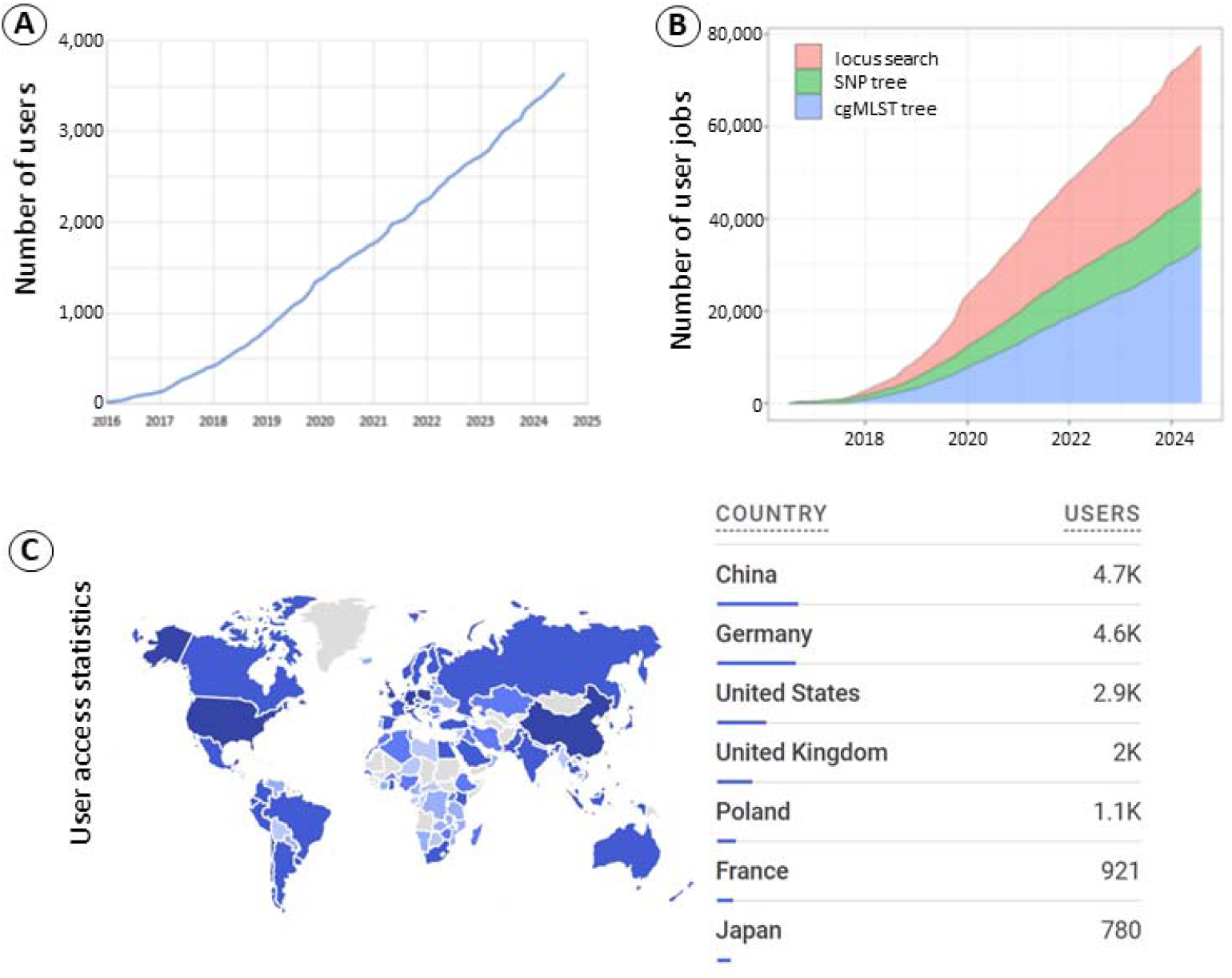
**(A)** Active EnteroBase users, 2016-2024. **(B)** Cumulative number of user jobs run on the EnteroBase platform. **(C)** The user access statistics by country as collected by Google Analytics for the period July 2023 to July 2024. The table shows the seven countries with the highest number of users.

## CONTENT

The total number of genome sequences in EnteroBase now exceeds one million, compared to 300,000 genomes in 2019 (Figure 2). This increase is partly due to the establishment of databases for additional pathogens, including *Streptococcus* (Warwick University site) and *Mycobacterium tuberculosis* (Leibniz Institute DSMZ site), which currently hold >120,000 genomes each. Most of this data was retrieved from public short-read archives, with automatic daily updates, whereas user uploads account for about 10% of the total data.

**Figure 2.**
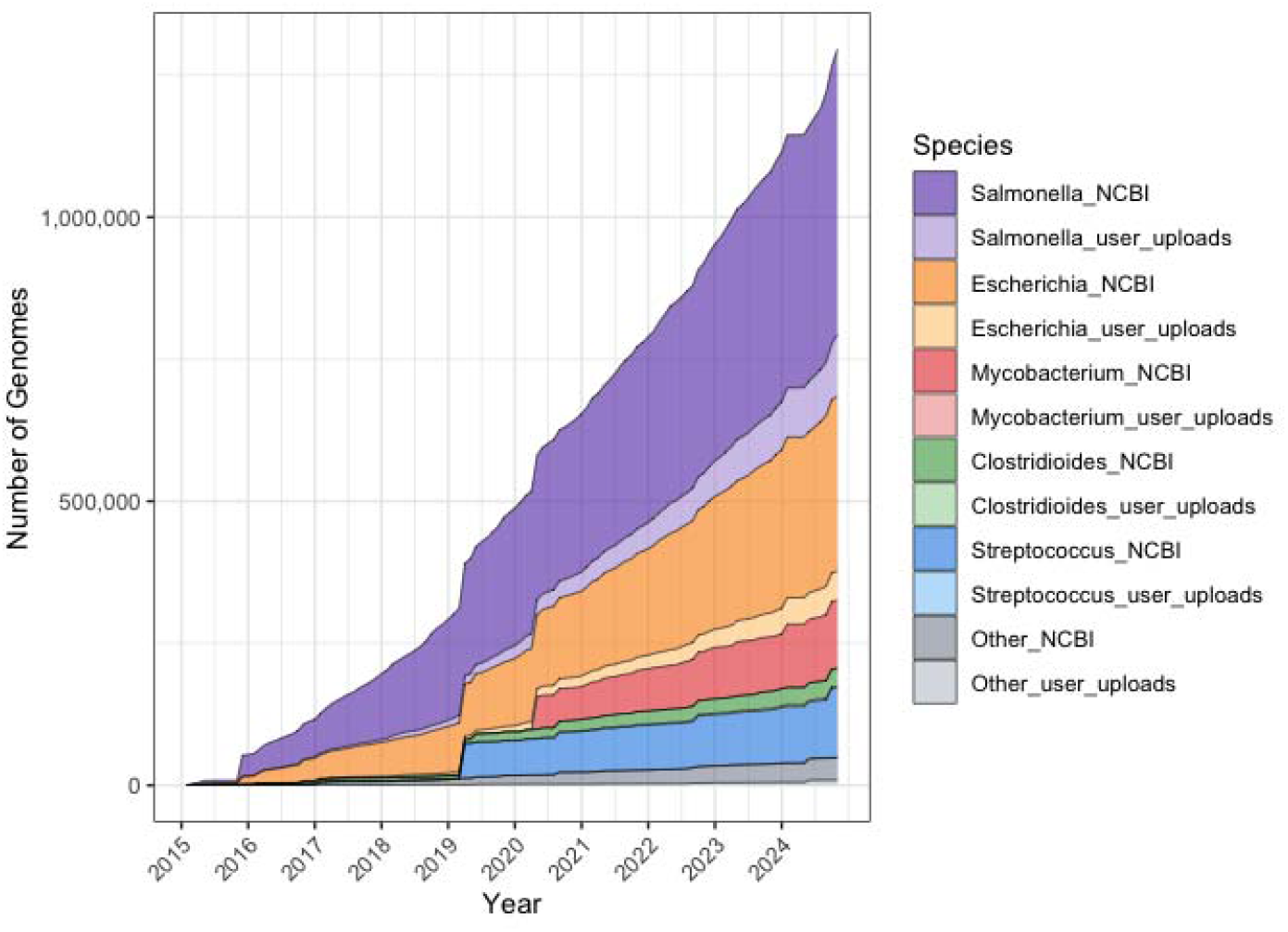
Increasing number of bacterial genome sequences provided by EnteroBase databases. The ‘other’ section includes the smaller databases (*Vibrio, Helicobacter, Moraxella and Yersinia*). Data sources are public short-read archives (NCBI) and user uploads, respectively.

The genome-based detection of antimicrobial resistance (AMR) determinants was implemented for *Salmonella, Escherichia* and *Clostridioides* databases and is currently indicated for each of 858,942 individual genomes (31 July 2024). Based on this new feature, AMR genotypes can be readily compared across tens of thousands of genomes to elucidate geographical distributions or temporal trends of specific resistance determinants, or their associations with specific sample types, disease patterns, and pathogen hosts (e.g. livestock vs. human). While other tools are available to predict AMR from genome sequences (9-11), they cannot easily compare AMR properties among large numbers of genomes, or across space and time.

We also found that the quality of AMR genotyping varied with the bacterial species under study and could be significantly improved by pathogen-specific data curation. For example, dedicated web tools failed to detect most genetic determinants conferring resistance against antibiotics recommended for the treatment of *Clostridioides difficile* infections, because associated databases were not up to date for this species (6). We have therefore recently implemented a modified pipeline for AMR detection for *C. difficile* in EnteroBase, together with an expert-curated database of genetic determinants to be detected (Figure 3). This pipeline detects plasmid sequences by using BLASTn (12) (version 2.2.31+; thresholds: sequence coverage ≥95%, sequence identity ≥99%), considering the circular nature of plasmids, and it detects mutations in promoter sequences by using BLASTn-short (thresholds: sequence coverage=100%, sequence identity ≥90%). In contrast, for the detection of point mutations in protein-coding genes (including base substitutions, insertions, and deletions), it relies on the AMRFinderPlus tool (9) (version 3.11.26) with an adapted database, screening for resistances using both protein and nucleotide reference sequences, respectively.

**Figure 3.**
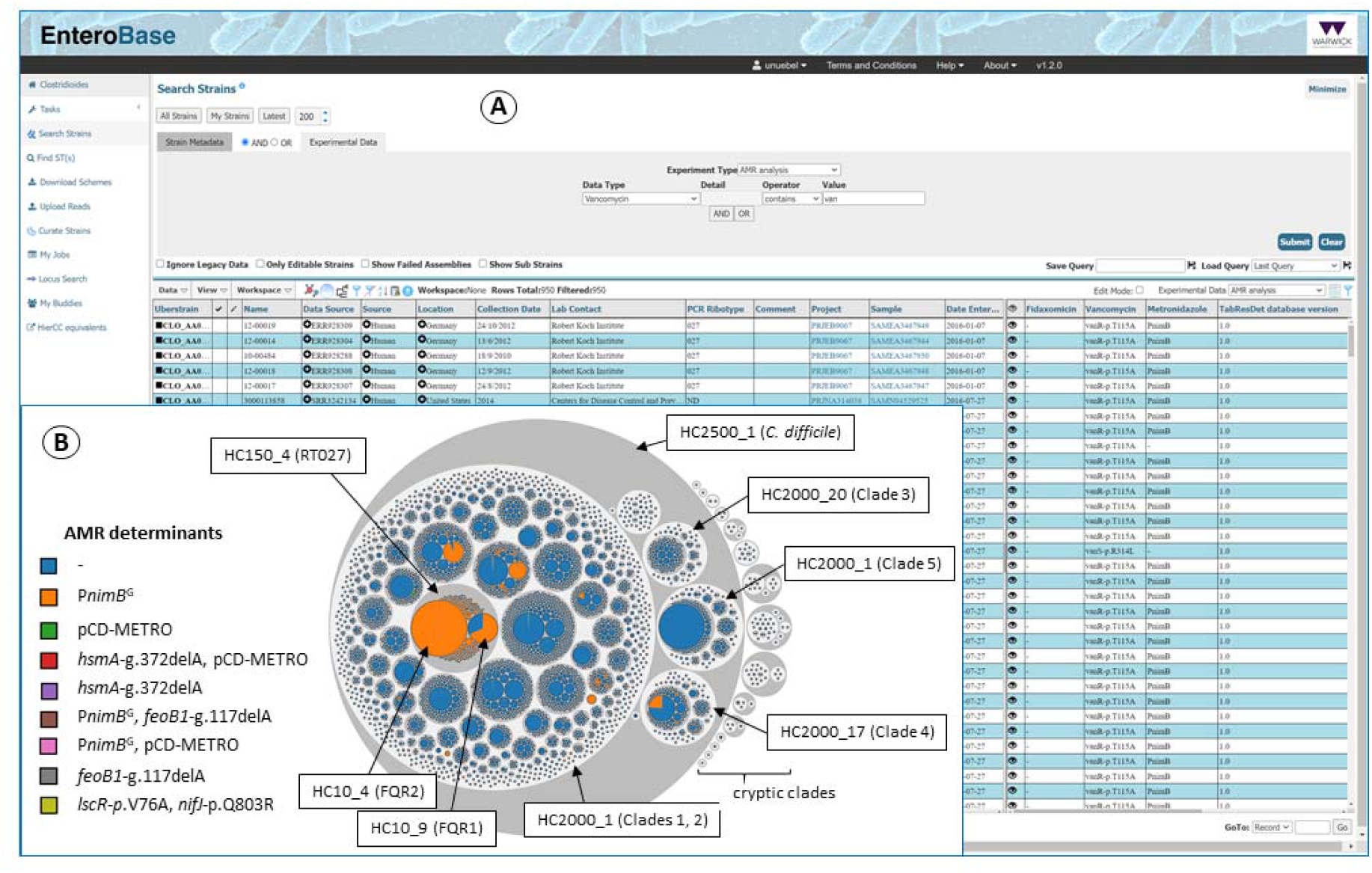
**(A)** *Clostridioides* genomes can be queried for specific genetic determinants for antimicrobial resistance using the Experiment Type ‘AMR analysis’. The returned genome entries are displayed in the spreadsheet. **(B)** Bubble plot illustrating the genetic population structure of *Clostridioides difficile* based on 30,599 genome sequences. All genomes are classified into hierarchical clusters at multiple levels, e.g. HC10 clusters identify chains of related genomes with pairwise differences of up to ten cgMLST alleles (see EnteroBase user guide/HierCC equivalents for correlates of hierarchical clusters in different bacterial species). Hierarchical clusters at levels HC150, HC2000 and HC2500, respectively, are indicated by grey shading, whereas HC10 clusters are colored based on the presence of genetic determinants of resistance to the antibiotic metronidazole, including plasmids and point mutations in both protein-coding genes and a promoter. It appears that the mutation in the *nimB* gene promoter (*PnimB*^G^), conferring increased metronidazole tolerance (13), is restricted to a limited number of genotypes, including epidemic PCR ribotype 027 strains FQR1 and FQR2 (correlating with hierarchical clusters HC10_9 and HC10_4, respectively (2)). Other determinants of metronidazole resistance, including the plasmid pCD-METRO, are rare in *C. difficile* (6). The interactive bubble plot shown here is available among public ‘workspaces’ within EnteroBase.

The AMR genotyping pipeline for *Salmonella* and *Escherichia* is also based on the AMRFinderPlus tool (9) (version 3.11.26). A transcription of the standard AMRFinderPlus output for each strain is available for viewing and download. On the GUI, AMR determinants are organised into columns by drug class. This mostly follows the AMRFinderPlus ‘class’, with the beta-lactam class further divided into penicillinases, extended-spectrum beta-lactamases (ESBL) and carbapenemases based on the ‘subclass’ categorization. Rare or clinically less relevant classes are grouped as ‘Other’ in the GUI. This distinction is important to identify remaining options for antibiotic treatment.

EnteroBase can now be used to search for bacterial strains with specific resistance determinants or classes, and AMR genotyping results can be displayed for a list of strains by selecting ‘AMR analysis’ from the ‘Experimental Data’ drop-down menu (Figure 3, Panel A). Additional details that may help to assess the confidence of the resistance genotype call are available by selecting the eye icon (e.g. the reference sequence type, length, and accession number, sequence coverage and identity, contig position, scientific literature).

## ANALYTICS TOOLS

We have implemented the new bubble plot graphical tool for visualising genetic population structures, based on pre-computed hierarchical clusters. In these plots, bubbles represent clusters of bacteria that are related at a specified level, with the bubble size indicating the number of genomes included, and they are each shown nested within the next-level hierarchical cluster. Users can choose to plot all levels of taxonomy, from clonal outbreaks to epidemics, and from endemic strains to the level of bacterial species. The bubble plots can be coloured using metadata or experimental data of interest to identify any correlations with population structure (Figure 3).

## USER INTERFACE

The graphical user interface of EnteroBase has been improved recently, with a unified design, interactive features and enhanced search functions. The EnteroBase websites are accessible by mobile devices, enabling genomic surveillance in the field, including rapid data synthesis and reporting. The homepage has been made more intuitive with descriptions of the main features to get a quick overview. All databases hosted by the University of Warwick and the Leibniz Institute DSMZ are displayed, indicating the genotyping schemes included and the total number of genomes available. Links have been added to the footer of the homepage for quick access to the user guide, contact information, terms and conditions, and acknowledgements. Additional pages provide information about the curators and developers involved in the maintenance and development of the EnteroBase software and databases, and on literature to be cited. For easy navigation, pages are titled and displayed pages are highlighted in the menu. The search functionality has been improved, allowing the user to view their query after search results have loaded. The query window is now integrated into the page (Figure 3) and automatically minimizes once a query has been submitted, to make room for the search results to be displayed. Registered users can save their search queries.

To enable integration with other platforms, EnteroBase now provides programmatic access to data and analysis tools via an application programming interface (RESTful API). It is designed for rapid live updates of small amounts of data. Queryable data includes cgMLST schemes, genome assemblies, genome metadata, locus sequences, allele profiles, and AMR predictions. All API activity must be requested via HTTP basic authentication and authenticated with a valid token. Requests can be wrapped in a programming language of the user’s choice; we recommend using Python and provide some example scripts at https://bitbucket.org/enterobase/enterobase-scripts. Responses are returned as JSON formatted data. To help users formulate their API requests, a Swagger sandbox is available at https://enterobase.warwick.ac.uk/api/v2.0/swagger-ui, which provides interactive documentation of the EnteroBase API, including information on endpoints, inputs, outputs and response codes.

## USER SUPPORT

The documentation of EnteroBase has been significantly improved and updated. Documentation on AMR analysis has been added and a guide to using API access to EnteroBase has been introduced, including several working examples. A wealth of detailed information for advanced users and developers has been added, to make it easier to install EnteroBase locally and to set up new databases and genotyping schemes.

Personal user support is provided through the support email address. The interactions with EnteroBase users continue to provide useful feedback which enables us to update the documentation and the tool itself to improve usability. New users often are guided to the specific documentation associated with the tasks they wish to perform. Occasionally, users will inform EnteroBase support of inconsistencies in the data held on EnteroBase, often arising from errors when data was originally uploaded to EnteroBase or NCBI. Such inconsistencies are then investigated and corrected. Users also ask about potential new features, which has helped set priorities for the adding of new features such as the introduction of AMR analysis.

## SYSTEM ARCHITECTURE

At the long-established Warwick University site, the EnteroBase software runs on four servers, with two servers each replicating each other’s functions to provide complete redundancy should one server fail. The PostgreSQL databases are mirrored in the same way. The four servers all reside on a secure network, with only a lightweight front-end webserver on the public internet. This machine serves some static html files, but its main role is to act as a load balancer, forwarding requests to the backend servers. The data is managed in a shared filesystem. In addition to the servers mentioned above there are further compute nodes that run the time and resource consuming jobs such as tree calculations and assemblies. At the Leibniz Institute DSMZ site, we are currently expanding the server structure to achieve a similar level of redundancy.

## FUTURE

EnteroBase will continue to be operated and actively developed at two locations. We are currently working on implementing AMR genotyping for additional species (e.g. *Mycobacterium tuberculosis*). We will establish databases for other pathogens and develop, install additional bioinformatics tools (e.g. for the identification of mobile genetic elements), and expand networking with other platforms through programmatic interfaces. EnteroBase will continue to be an important resource for making efficient use of the increasing amount of pathogen sequencing data and realising the full potential of high-throughput genome sequencing.

## ACKNOWLEDGEMENTS

The EnteroBase system was originally developed by Prof. Mark Achtman and his research group at the University of Warwick. We are most grateful to our expert database curators, including M. Chattaway, M. Pardos, F.-X. Weill, E. Litrup, S. Beatson, C. Nodari, M. Kilian, M. Frentrup, B. Gomez-Gil, C. Constantinidou, K. Thorell, R. Torres, S. Reuter, S. Nguyen. The authors at Leibniz Institute DSMZ acknowledge continuous support from their scientific computing team. The graphical abstract was created using freely available images from Flaticon.com.

